# De novo transcriptome assembly, functional annotation and characterization of the Atlantic bluefin tuna (*Thunnus thynnus*) larval stage

**DOI:** 10.1101/2020.05.27.118927

**Authors:** Marisaldi Luca, Basili Danilo, Gioacchini Giorgia, Carnevali Oliana

**Affiliations:** Department of Life and Environmental Sciences, Università Politecnica delle Marche, Ancona 60131, Italy; Centre for Molecular Informatics, Department of Chemistry, University of Cambridge, Lensfield Road, Cambridge, CB2 1EW, UK

**Keywords:** Larval transcriptome, *Thunnus thynnus*, tuna larvae

## Abstract

Over the last two decades, many efforts have been invested in attempting to close the life cycle of the iconic Atlantic bluefin tuna (*Thunnus thynnus*) and develop a true aquaculture-based market. However, the limited molecular resources nowadays available represent a clear limitation towards the domestication of this species. To fill such a gap of knowledge, we assembled and characterized a de novo larval transcriptome by taking advantage of publicly available databases with the final goal of better understanding the larval development. The assembled transcriptome comprised 37,117 protein-coding transcripts, of which 13,633 full-length (>80% coverage), with an Ex90N50 of 3,061 bp and 76% of complete and single-copy core vertebrate genes orthologues. Of these transcripts, 34,980 had a hit against the EggNOG database and 14,983 with the KAAS annotation server. By comparing our data with a set of representative fish species proteomes, it was found that 78.4% of the tuna transcripts were successfully included in orthologous groups. Codon usage bias was identified for processes such as translation, peptide biosynthesis, muscle development and ion transport, supporting the idea of mechanisms at play in regulating stability and translation efficiency of transcripts belonging to key biological processes during the larval growth. The information generated by this study on the Atlantic bluefin tuna represent a relevant improvement of the transcriptomic resources available to the scientific community and lays the foundation for future works aimed at exploring in greater detail physiological responses at molecular level in different larval stages.

## 1 Introduction

The Atlantic bluefin tuna Thunnus thynnus is a well-known iconic species and one of the biggest fish in the pelagic realm. Due to a high geographical distribution range, exceptional migratory behaviour and high commercial interest, the management of the fishing activities requires coordinated international cooperation and networks, a task managed by International Commission for the conservation of Atlantic Tunas (ICCAT). As a result of this international conservation effort to preserve the species, the Atlantic population is divided into two management units, an Eastern and Western stock. However, an ever-increasing demand over recent years, mainly driven by the sushi and sashimi Japanese market, has led the Eastern stock to the brink of collapse (MacKenzie et al. 2009). Even if harsh, the restrictive measures put in place (ICCAT 2008) were effective to recover the stock but the risk taken also suggested that the time was ripe to develop an aquaculture-based market.

Any large-scale aquaculture operation requires full control of the production cycle of the species of interest, a key point not yet accomplished for the Atlantic bluefin tuna. As a matter of fact, current practices to raise this species in captivity consist of catching large schools of wild fish, transport and feed them at sea-based floating cages for several months to gain weight and obtain the desired fat composition to meet the market demand (Mylonas et al. 2010). Such a production model is heavily dependent on the fishing activities and as a consequence, on the natural fluctuations of the wild population.

One of the first point to be addressed to achieve the domestication is a thorough knowledge of the reproductive cycle and the captivity effects on gametogenesis and endocrine axis (Medina et al. 2016; Zohar et al. 2016; Carnevali et al. 2019). Recently, more attempts to manipulate the life cycle (i.e. induce spawning) and grow the larvae were carried out with promising achievements (Mylonas et al. 2007; De Metrio et al. 2010; Reglero et al. 2014; Yúfera et al. 2014; Blanco et al. 2017; Betancor et al. 2017; Betancor et al. 2019; Betancor et al. 2020). In this context, understanding nutritional requirements, growth pattern, responses to external stimuli, morphological and physiological changes and discovering stage-specific markers during the larval development are central points (Gisbert et al. 2008; Rønnestad et al. 2013). A deep knowledge of the genes driving these processes is therefore necessary to obtain a more complete picture. However, one of the constrains to figure out requirements and best farming conditions for this species is a surprising lack of molecular information, from genomic and transcriptomic resources to simple availability of sequences in databases. As a rough comparison, for the Pacific bluefin tuna *Thunnus orientalis*, the completion of the life cycle was achieved back in 2002 (Sawada et al. 2005) and a recent work by Suda et al. (2019) improved the first version of the genome by discovering male-specific DNA markers useful for the control of the sex ratio in tuna farms.

Filling such a gap of knowledge would represent an important step to support ongoing efforts towards the domestication of the Atlantic bluefin tuna and to boost the research in this direction. For these reasons, by taking advantage of publicly available databases, the aim of the present study was to generate, annotate and characterize a de novo Atlantic bluefin tuna larval transcriptome assembly from a comprehensive pool of several larval stages. Orthologous groups with other fish species were established by means of a comparative analysis and, along with the analysis of codon bias, allowed for the identification of important molecular processes during larval ontogeny. The information generated by this study, through the identification of the main features of the larval transcriptome, represent a significant advance of the transcriptomic resources available for this species and will support future studies aimed at investigating physiological responses at molecular level in different larval stages.

## 2 Material and methods

### 2.1 Data availability, pre-processing steps and de novo assembly

The experimental dataset (100 bp paired-end RNA-seq) from a pool of larvae at different developmental stages (0dph, 2dph, 10dph, 15dhp, 20dph) was downloaded from the NCBI Sequence Read Archive (SRA; Accession number: SRR1536893) and analysed with a rigorous bioinformatic workflow (Fig. 1). For this raw data, neither a publication nor any report and/or related information was found at the time of the download (December 2019). The quality of the raw reads was first assessed with FastQC (https://www.bioinformatics.babraham.ac.uk/projects/fastqc/). Along the workflow, FastQC was used before and after each step to compare the benefits derived from each tool applied (Fig. 1). First, ribosomal RNA was filtered out with SortMeRna (Kopylova et al. 2012) by screening the raw reads against the provided representative rRNA databases. Then, potential bacterial, archaeal and viral contamination was eliminated with Kraken2 (Wood et al. 2019) using the standard Kraken database. Decontaminated reads were further processed with Prinseq (Schmieder and Edwards 2011) to remove low complexity sequences and trim long poly-A/T tails. As a last step before the assembly process, a gentle trimming was applied with Trimmomatic (Bolger et al. 2014) with parameters adjusted according to MacManes (2014). The resulting cleaned reads were used for the de novo assembly step with Trinity (Haas et al. 2013) without the in silico normalization. The level of strand-specificity of the library was checked with the Trinity script *examine_strand_specificity.pl*. The complete list of specific parameters set for each software with the relative version is available in Supplementary file S1.

**Figure 1:**
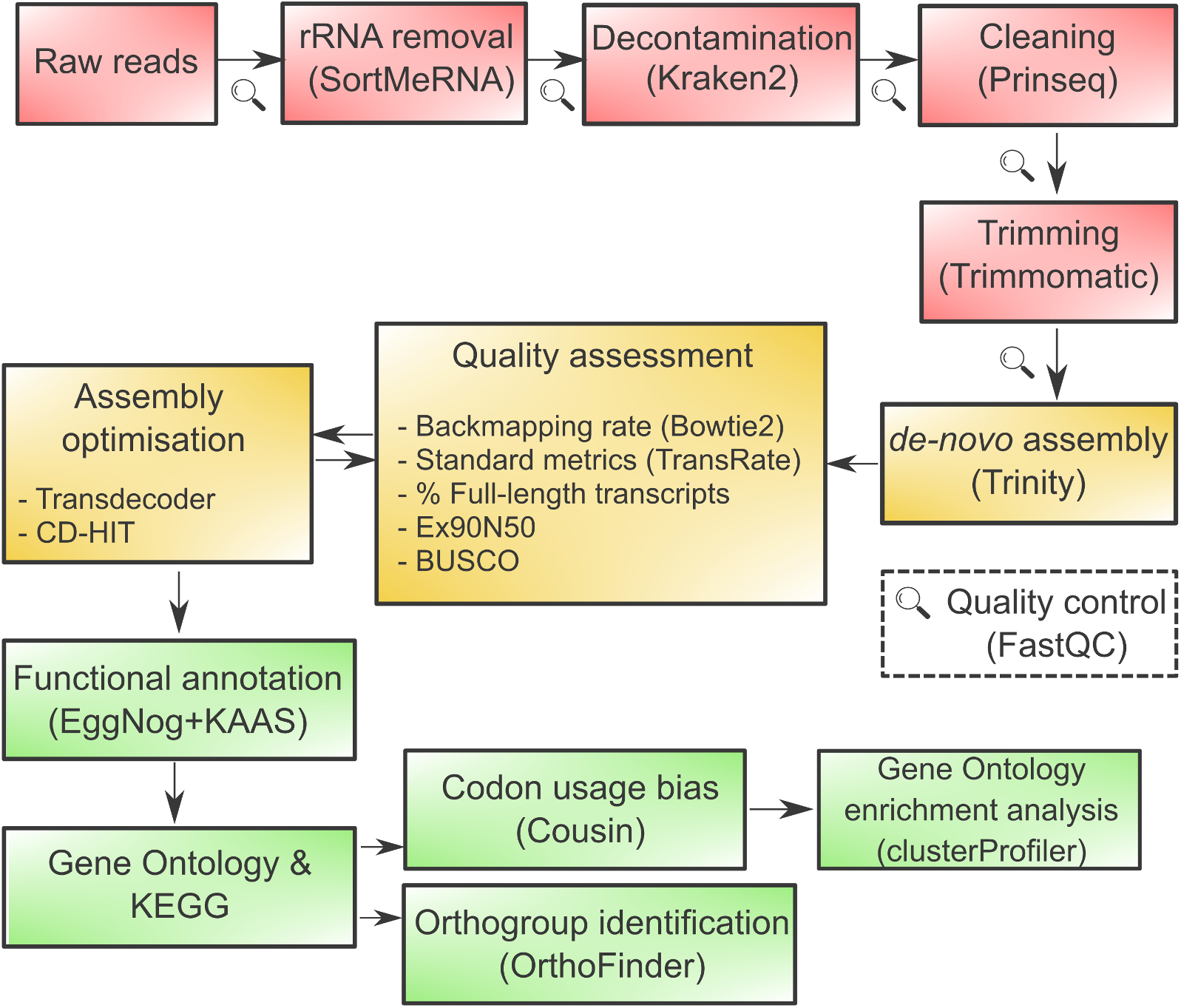
Schematic workflow of the analysis carried out to generate and annotate the de novo transcriptome assembly of the Atlantic bluefin tuna larvae. The tools used at each step are reported in brackets.

### 2.2 Assembly optimisation and functional annotation

Cleaned reads were mapped back to the raw transcriptome assembly by applying Bowtie2 (Langmead and Salzberg 2012) and the overall metrics were calculated with Transrate (Smith-Unna et al. 2016). Then, the completeness of the assembly was assessed with BUSCO (Simão et al. 2015) using the Actinopterygii odb9 database and gVolante (Nishimura et al. 2017) using a set of Core Vertebrate Genes (CVG) as identified by Hara et al. (2015). The percentage of full-length reconstructed transcripts was calculated with the *analyze_blastPlus_topHit_coverage.pl* script of the Trinity package by blasting the translated transcriptome against the proteome (Sedor1 version) of the Yellowtail amberjack (*Seriola lalandi dorsalis*) downloaded from Ensembl release 98 (www.ensembl.org). The rationale behind that is the availability of the complete genome for this species and the relatively close phylogenetic relationship of the Carangiformes with tunas (Hughes et al. 2018). As an additional assessment, the N50 metric was integrated with the expression data computed with the Trinity tool suite (*align_and_estimate_abundance.pl; abundance_estimates_to_matrix.pl*) using Kallisto (Bray et al. 2016) as quantification method. This produced the Ex90N50 metric, that is the N50 statistic limited to the highly expressed transcripts that represent 90% of the total normalized expression data (for detailed methods see the Trinity GitHub page at https://github.com/trinityrnaseq/trinityrnaseq/wiki/Transcriptome-Contig-Nx-and-ExN50-stats).

The raw transcriptome was then optimised with TransDecoder (Haas et al. 2013) by keeping the single best Open Reading Frame (ORF) per transcript and with CD-HIT (Fu et al. 2012) to cluster transcripts with 100% aa identity. To assess the quality gained with the optimisation procedure, the same checkpoints adopted for the raw transcriptome were re-applied and the corresponding results compared between the two versions (Fig. 1). Most of the steps required up to the quality assessment were performed on the public Galaxy-Europe platform (Afgan et al. 2018). Once optimised, the transcriptome was annotated with EggNog-mapper v2 (Huerta-Cepas et al. 2017) and the Gene Ontology terms analysed with CateGorizer using the GO slim terms as reference (Zhi-Liang et al. 2008). Furthermore, the transcriptome was also annotated with the KEGG Automatic Annotation Server (KAAS) using the Blast bi-directional best hit (BBH) method (Moriya et al. 2007). The results of this step were then analysed with the “KEGG mapper-search and colour pathways” using “reference” as search mode. The information and number of genes/pathway were retrieved using KEGGREST (Tenenbaum 2018).

### 2.3 Codon usage bias analysis

Codon usage bias was assessed by calculating the Effective Number of Codons (ENCs), a measure of how much codon usage preferences are distant from an equal usage of synonymous codons (Wright 1990). According to the Transdecoder output, only the coding sequences from transcripts with a complete open reading frame were retained for this analysis. The calculation of the ENCs was performed with COUSIN (Bourret et al. 2019). The transcripts with an ENCs value falling below the first quartile of the distribution were considered as having stronger codon usage bias and therefore isolated for downstream analysis. Then, the Gene Ontology enrichment analysis was performed for these transcripts with clusterProfiler (Yu et al. 2012) in the RStudio environment (RStudio team 2015). The *p*-values (cut-off 0.05) calculated by the hypergeometric distribution were adjusted with the BenjaminiHochberg method (Benjamini and Hochberg 1995). The redundancy of terms from the enrichment results was filtered with Revigo (Supek et al. 2011) using default parameters.

### 2.4 Comparative analysis

The optimised transcriptome was compared with the proteome of other fish species, for which a well-annotated genome was available on Ensembl release 98, by using OrthoFinder to identify orthologous groups, also called orthogroups (Emms and Kelly 2015). As explained by the authors, an orthogroup is the extension of the notion of orthology to groups of species, meaning that an orthogroup is the group of genes descended from a single gene in the last common ancestor of a group of species. For this analysis, the chosen species were the zebrafish (*Danio rerio*), the spotted gar (*Lepisosteus oculatus*), the yellowtail amberjack (*Seriola lalandi*), the Nile tilapia (*Oreochromis niloticus*), the sea bream (*Sparus aurata*) and the pufferfish (*Tetraodon nigroviridis*). The species were chosen to be representative of broad taxonomical groups of fish. To visualise intersections between groups, the R package UpSetR (Conway et al. 2017) was chosen instead of the Venn diagrams due to the relatively high number of intersections. The phylogenetic tree from the OrthoFinder analysis was plotted with the online tool iTOL (Letunic and Bork 2007). Then, “assembly-specific” transcripts included only in tuna groups by OrthoFinder were further screened by using Revigo (Supek et al. 2011), with a small allowed similarity (C=0.5) and “SimRel” as semantic similarity to explore the associated Gene Ontology terms.

In addition to the aforementioned tools, the final plots were created in the RStudio environment with the following packages: ggplot2 (Wickham 2016), ggpubr (Kassambara 2020) and ggrepel (Slowikowski 2020). The black silhouettes used for the final figures were downloaded from the PhyloPic website (http://phylopic.org/).

## 3 Results

### 3.1 Transcriptome assembly and quality assessment

The raw data were composed by 29,910,574 millions of Illumina paired-end reads which were narrowed to 27,379,333 after the filtering and trimming steps. A comparison between the metrics and quality of the raw and optimised transcriptomes is summarised in Table 1. Up to 65.5% of the transcripts were dropped in the optimised transcriptome without significantly affecting the overall quality, suggesting that a consistent fraction of the reconstructed transcripts was background noise derived from lowly expressed transcripts. Raw transcriptome benchmarking against the Actinopterygii single-copy orthologues database (odb9, n=4584) found 88.2% complete (Single:52.0%, Duplicated:36.2%), 6.2% fragmented and 5.6% missing BUSCOs while 87.8% complete (Single:70.3%, Duplicated:17.5%), 6% fragmented and 6.2% missing BUSCOs in the optimised transcriptome (Fig. 2A). Similar results were obtained by benchmarking the two transcriptomes against the CVG database (n=233). Indeed, for the raw version it was found 90.6% complete (Single:56.7%, Duplicated:33.9%), 6.9% fragmented and 2.5% missing BUSCOs while 90.2% complete (Single:76.0%, Duplicated:14.2%), 6.0% fragmented and 3.8% missing BUSCOs for the optimised version (Fig. 2A). The shape of the Ex90N50 curves for the initial and optimised assembly confirmed the efficacy of the post-assembly optimising procedure (Fig. 2B) as outlined by the developers on the Trinity wiki page (https://github.com/trinityrnaseq/trinityrnaseq/wiki). The analysis of the strandness revealed non-strand-specific RNA-Seq data. Since the quality assessment revealed a good improvement following the optimising steps, all the downstream analyses were carried out on the optimised transcriptome. The optimised transcriptome is available as Supplementary material S2.

**Table 1:** Statistics regarding the quality assessment of the raw and optimised transcriptomes.

| Statistics | Raw | Optimised |
| --- | --- | --- |
| N sequences | 107,465 | 37,117 |
| GC content | 45% | 48% |
| N50 | 2835 bp | 3267 bp |
| Ex90N50 | 2774 bp | 3061 bp |
| Backmapping rate | 98.30% | 83.71% |
| N full-length transcripts (>80% coverage) | 13,972 (13%) | 13,633 (36%) |

**Figure 2:**
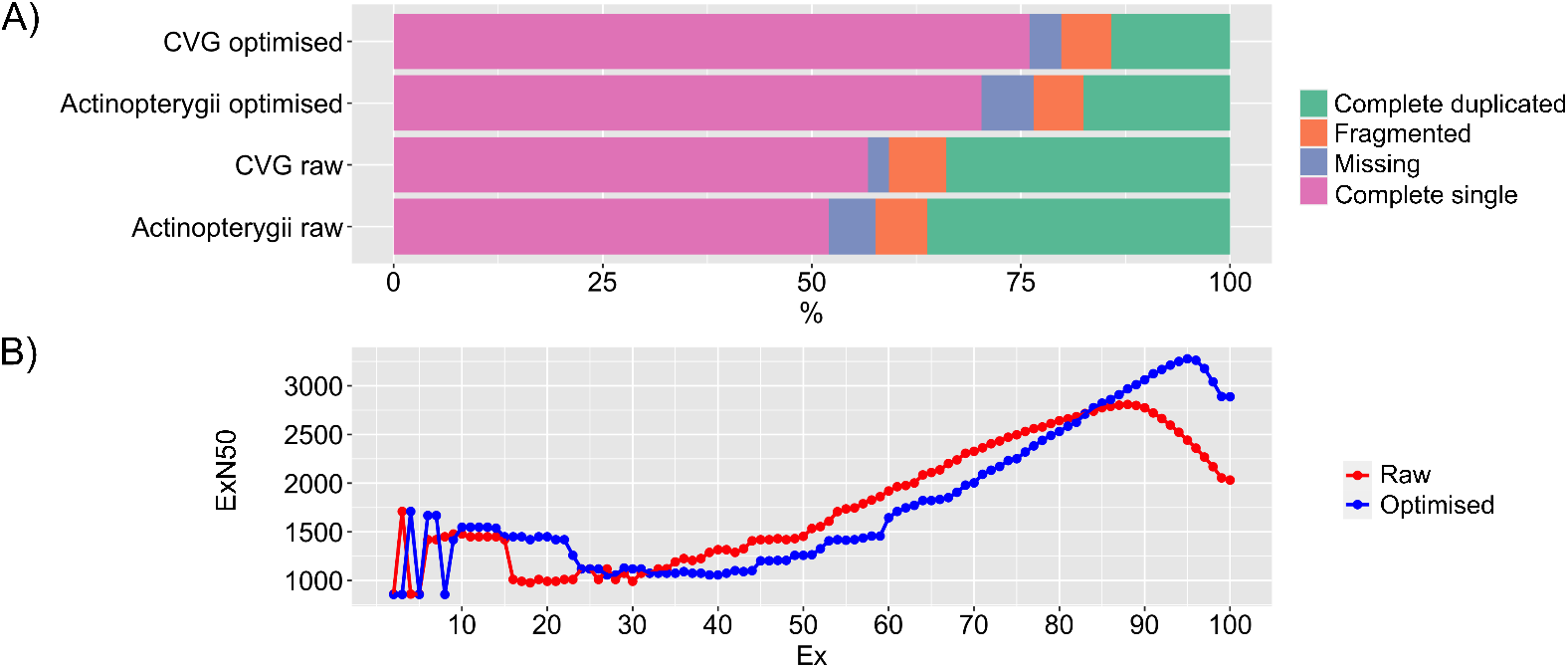
Quality assessment between the raw and optimised transcriptomes A) percentages of BUSCOs identified when searched against the Actinoptery-gii database (odb9) and the set of Core Vertebrate Genes (CVG); B) ExN50 curves representing the percentage of the total normalized expression data (x-axis) with the relative N50 values (y-axis).

### 3.2 Functional annotation

A total of 34,980 transcripts (94.2% of the total sequences) had a hit against the EggNOG database, of which 25,539 (72.99%), 26,024 (74.38%) and 28,836 (77.6%) resulted associated with the relative GO terms, KEGG orthology functional annotation and gene symbols, respectively (Supplementary file S3). The KAAS annotation server assigned a KEGG orthology identifier to 14,983 transcripts (Supplementary file S3). The GO slim terms by functional category (biological process, molecular function and cellular component) are summarised in Fig. 3. The most represented biological processes were related to development (GO:0008152), metabolism (GO:0007275), cell organization (GO:0016043), transport (GO:0006810) and cell differentiation (GO:0030154). The terms intracellular (GO:0005623), cytoplasm (GO:0005622) and cell (GO:0005737) were the top represented cellular components. Moreover, among the molecular functions there were catalytic (GO:0003824), hydrolase activity (GO:0016787), transferase activity (GO:0005488) as well as binding (GO:0016740). Among the most interesting KEGG pathways with a relevant number of mapped genes there were metabolism-related pathways (fatty acids elongation: map00062; biosynthesis of unsaturated amino acids: map01040, arachidonic acid metabolism: map00590, PPAR signalling: map03320), phototransduction (map04744), GnRH signalling (map04912), thyroid hormone signalling (map04919) and cell death-related pathways (autophagy-animal: map04140, apoptosis: map04210) (Fig. 3).

**Figure 3:**
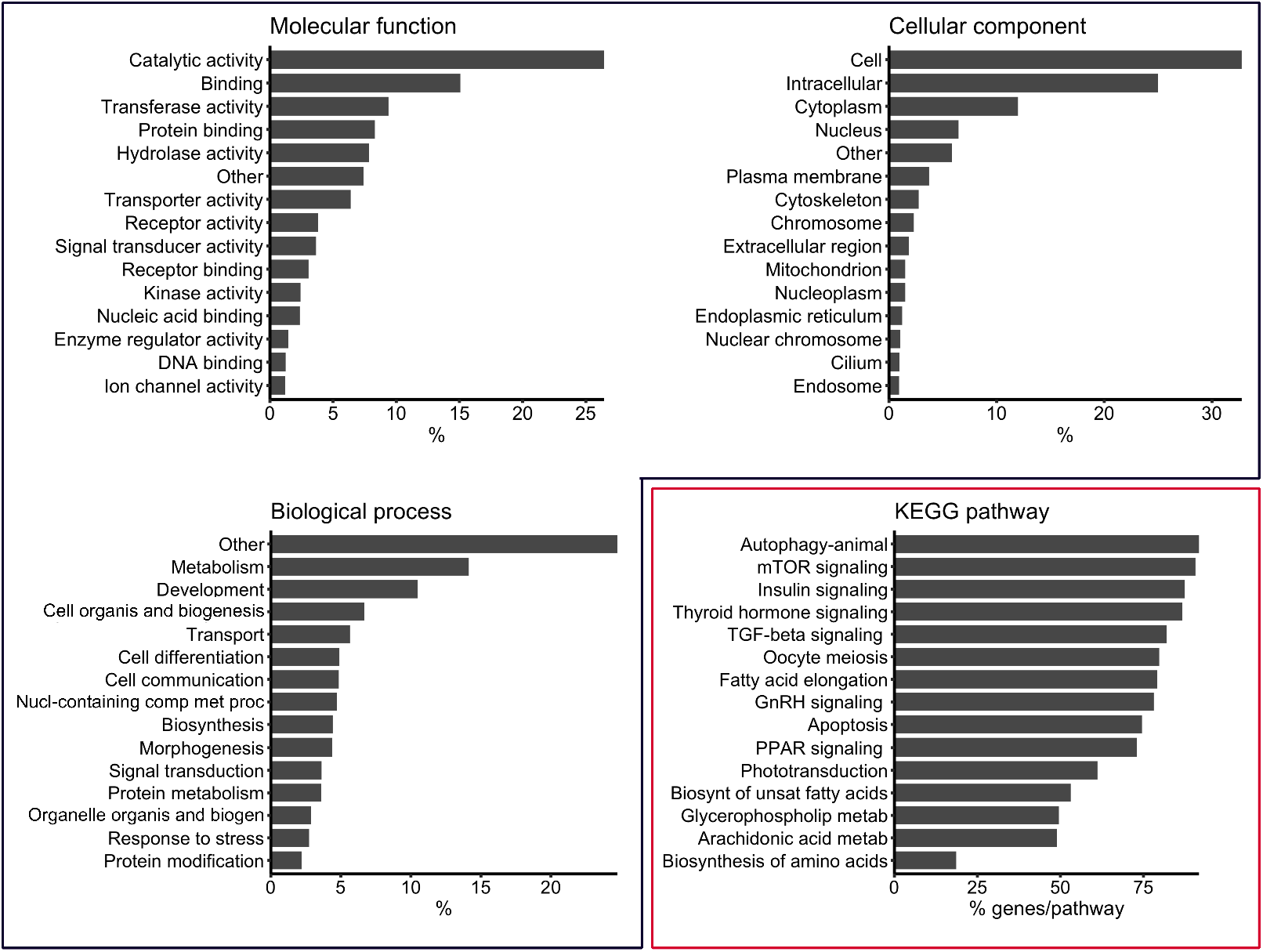
Proportion of Gene Ontology annotations for the three categories corresponding to molecular functions, cellular component and biological process (black box) and the number of genes identified in relevant KEGG path-ways (red box).

### 3.3 Selection of codons during the larval development

The distribution of the ENCs was slightly skewed to lower values with a mean value of 49.88 and the first quartile of 47.31. A total of 3667 transcripts were found below the first quartile of the ENCs distribution (i.e. more biased codon usage) and used for the enrichment analysis. Accordingly, some of the statistically significant over-represented biological processes were related to translation (GO:0006412), muscle organ development (GO:0007517) and microvillus assembly (GO:0030033). Among the cellular component, there were ribosome-related terms (ribosomal subunit: GO:0044391; large ribosomal subunit: GO:0015934; cytosolic ribosome: GO:0022626), sarcomere (GO:0030017), myelin sheath (GO:0043209) and presynapse (GO:0098793). Instead, among the molecular functions, there were terms linked to ion transmembrane transport (ATPase activity, coupled to transmembrane movement of ions: GO:0044769, active ion transmembrane transporter activity: GO:0022853, ATPase-coupled ion transmembrane transporter activity: GO:0042625). The results of the Gene Ontology enrichment analysis are reported in Fig. 4. The complete list of both raw and filtered enriched terms can be found in the Supplementary file S4.

**Figure 4:**
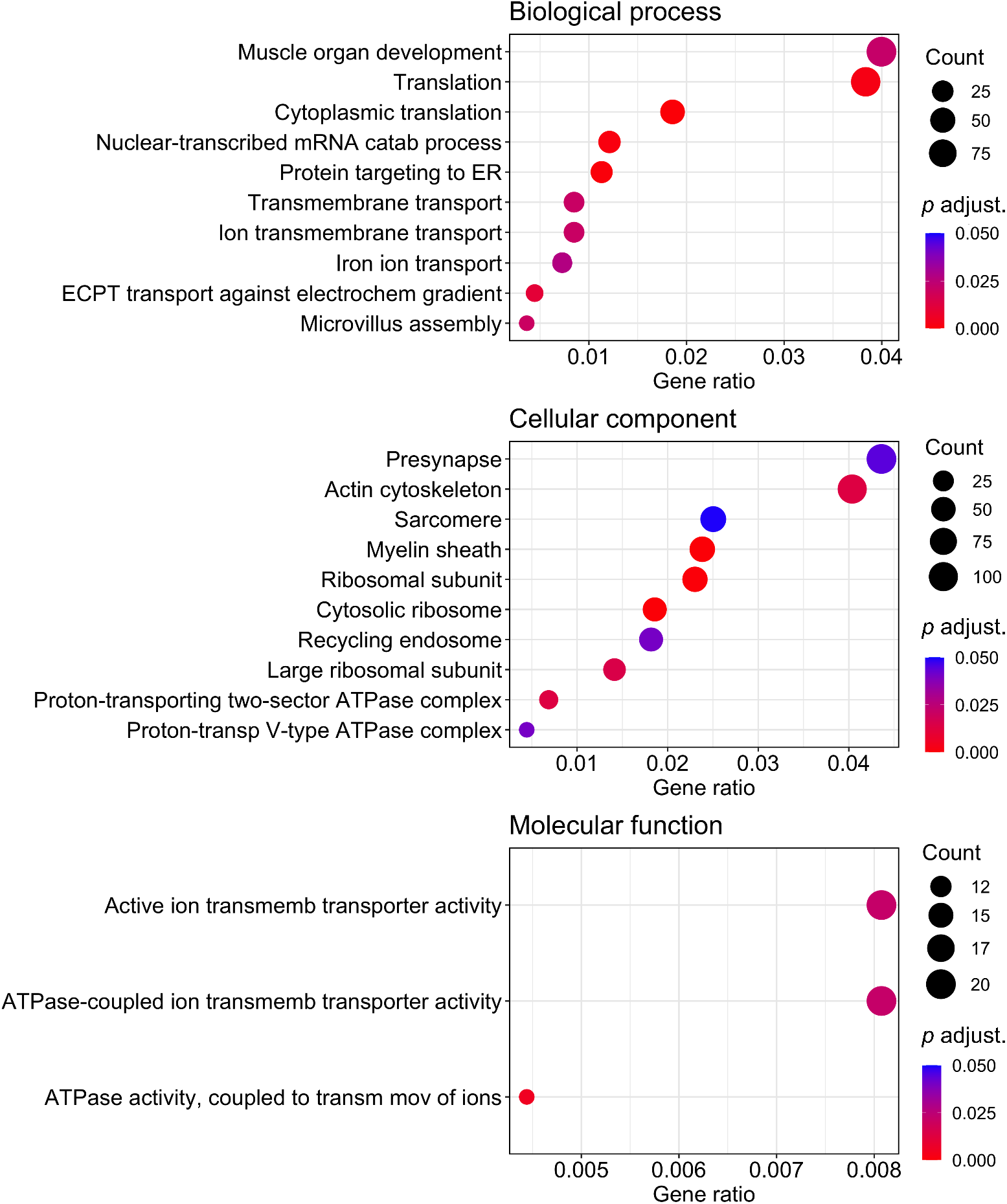
Gene Ontology enrichment analysis of transcripts falling below the first quartile of the ENCs distribution corresponding to lower values (i.e. more codon usage bias). The gene ratio (x-axis) corresponds to k/n in which k is the number of annotated genes in the gene list of interest for a particular term and n is the size of the list of gene of interest. Abbreviations: ECPT: energy coupled proton transmembrane; ER: endoplasmic reticulum.

### 3.4 Orthogroups identification

The OrthoFinder analysis identified 10,995 orthogroups with all the species present (Fig. 5A) and a tiny fraction of genes was in species-specific groups ranging between 0.1 to 1.1% of the total (Supplementary file S5). Between 78.4% and 93.9% of the genes across the seven species were assigned to orthogroups and between 6.1%-21.6% of the total was not assigned to any orthogroup (Supplementary file S5). A total of 740 single-copy orthogroups, which corresponds to groups containing exactly 1:1 orthologue proteins, were identified. The complete and single-copy list of orthogroups can be retrieved from the Supplementary material S6. The tuna transcriptome resulted in 29,090 (78.4%) and 8,027 (21.6%) transcripts assigned and unassigned to orthogroups, respectively. A total of 16 “assembly-specific” tuna groups (Fig. 5B), containing 64 transcripts, were found. Of these transcripts, 50 were associated with RefSeq, Ensembl or TrEMBL IDs but as low as 14 were well characterized and assigned also to GO terms and 23 to KEGG orthologues according to the EggNog annotation. Interestingly, several “assembly-specific” transcripts belonged to the chymotrypsin family (peptidase S1A), a group of enzymes with roles in protein digestion and absorption as well as pancreatic secretion. A manual inspection showed that such transcripts had a complete coding sequence, well defined protein domains and family classification as revealed by InterProScan (http://www.ebi.ac.uk/interpro/) and a low identity (~30%) with other fish species according to NCBI blast results. In general, among the interesting biological processes associated with the “assembly specific” transcripts, there was iris morphogenesis (GO:0061072), regulation of smoothened signalling pathway (GO:0008589), negative regulation of wound healing (GO:0061045), actin-mediated cell contraction (GO:0070252) and the regulation of fat cell differentiation (GO:0045598) (Fig. 6). According to the Revigo output, the biological process and cellular component terms were filtered for those representative clusters including more than 5 terms to reduce overplotting (see Supek et al., 2011 for details). The complete list of transcripts belonging to these “assembly-specific” tuna groups as well as the full results of Revigo can be found in the Supplementary material S7. The phylogenetic relationships between the selected species, reconstructed by OrthoFinder, are displayed in Fig. 5C (see Emms and Kelly (2019) for details about the methods).

**Figure 5:**
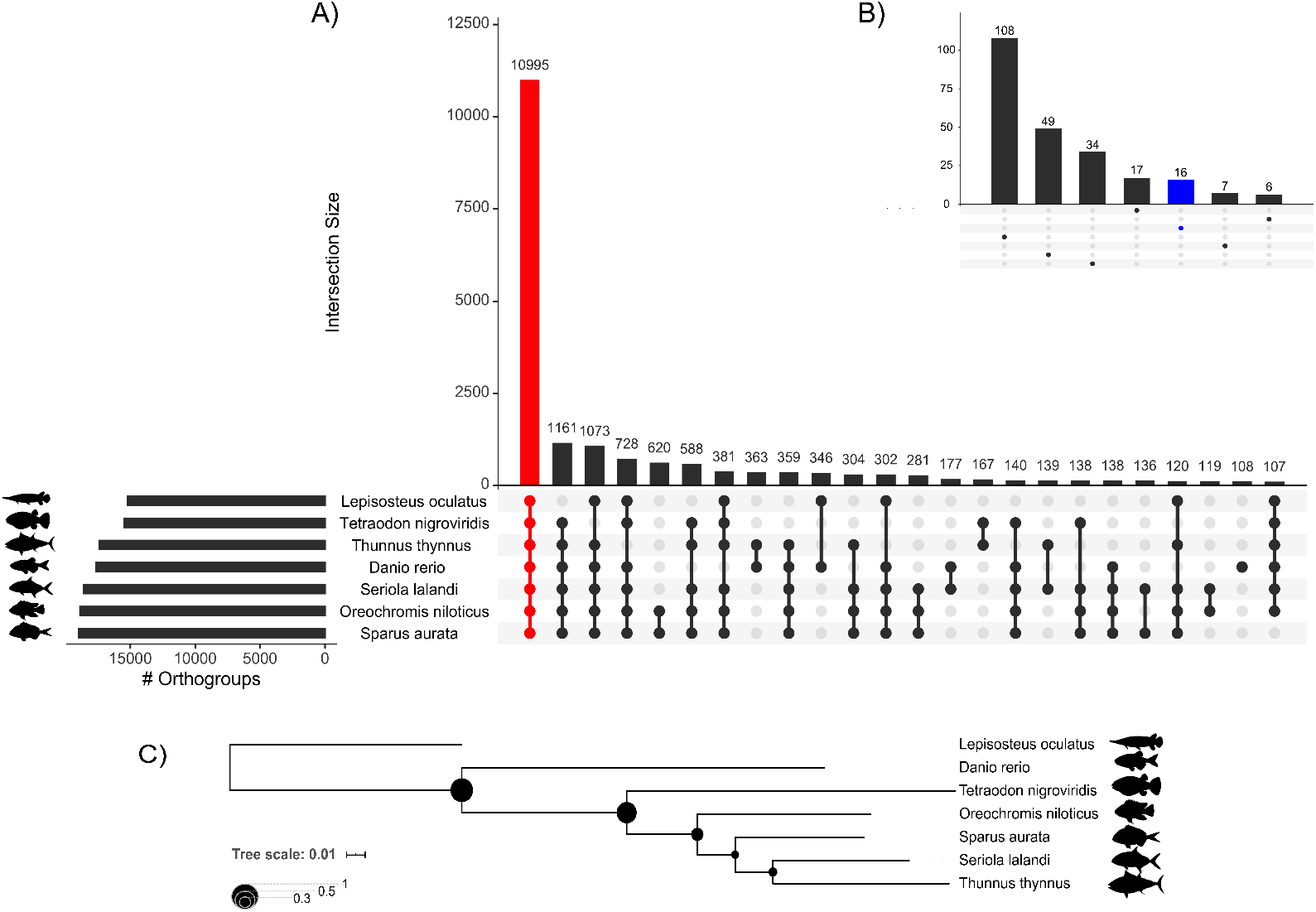
Number of genes in orthogroups in the selected species (A) and in species-specific groups (B). The red bar corresponds to orthogroups shared by all the species considered in the analysis and the blue bar corresponds to the 16 “assembly-specific” tuna groups. Numbers above each bar represent the number of orthogroups shared in that particular intersection between species. C) The phylogenetic tree reconstructed by the OrthoFinder analysis.

**Figure 6:**
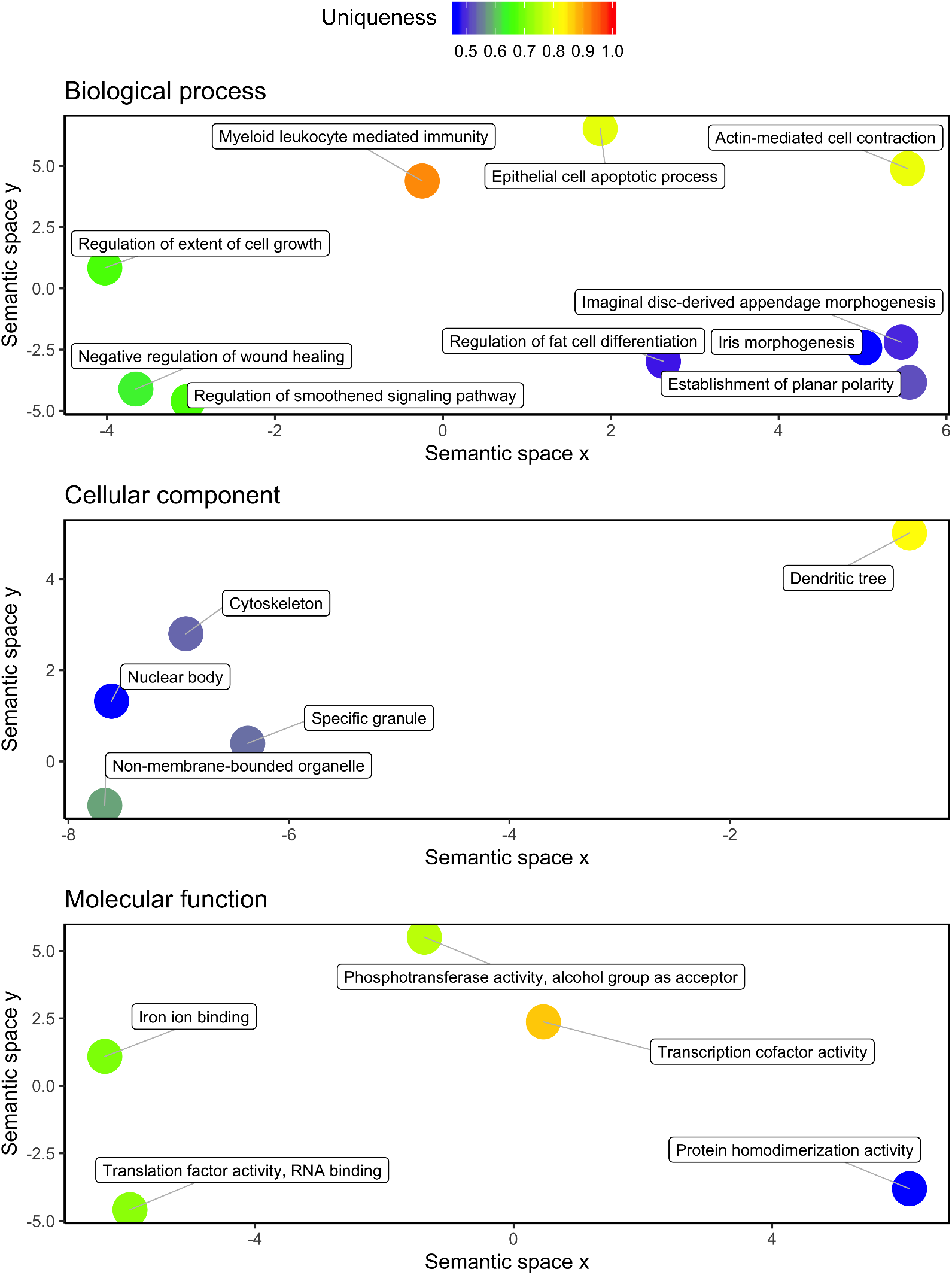
Gene ontology terms associated with “assembly-specific” transcripts in the Atlantic bluefin tuna plotted using the Revigo results. The distance between GO terms on the plot is proportional to their semantic similarity according to the principles and methods outlined by Supek et al. (2011).

## 4 Discussion

The high taxonomical diversity of teleost fishes is often accompanied by astonishing physiological and morphological changes during the larval metamorphosis, giving to a single species the advantage of exploiting different habitats and ecological niches (McMenamin and Parichy 2013). For this reason, raising any target species in captivity requires a thorough understanding of both the mechanisms driving the transition from the larval to the adult stage as well as its biology, ecology and nutritional needs. Therefore, the identification and characterization of the genes and molecular pathways driving traits of commercial interests such as growth rates, reproduction, larval development and immune response would implement production and management models in the industry. The efforts in this direction are becoming routinely supported by transcriptomic approaches in both promising new farmed species as the Atlantic bonito (Sarropoulou et al. 2014) and well established farmed fish as salmonids (Carruthers et al. 2018) and soles (Benzekri et al. 2014). In many cases, the transcriptomic resources paved the way to genome projects due to remarkable technological progress and plummeting sequencing costs, making a genome assembly affordable by numerous research groups. This was the timeline for species of key commercial interest such as the sea bream (Calduch-Giner et al. 2013; Pauletto et al. 2018), the European sea bass (Louro et al. 2010; Tine et al. 2014) and flatfish (Cerdà and Manchado 2013). In this context, the first study in the Atlantic bluefin tuna with similar efforts dates back more than a decade ago with the application of expressed sequence tag (EST) technology to investigate expression libraries in ovary, testis and liver (Chini et al. 2008). Then, Trumbić et al. (2015) applied RNA-seq on Atlantic bluefin tuna with the main purpose of developing a DNA microarray, therefore not even a single transcriptome was publicly available for this species, despite being a hot topic in the field of aquaculture. Whilst the central point is a matter of research efforts, this lack of resources was also linked to the unavailability of assembled transcripts and annotations along with relatively published papers. It follows that this de novo larval transcriptome assembly represents a first crucial step towards the domestication of this species and a significant progress of the genomic resources available for tunas.

According to general principles which start to be quite well established after more than a decade of de novo transcriptomes (Delhomme et al. 2014; Conesa et al. 2016; Moreton et al. 2016), the parameters chosen to assess the tuna transcriptome revealed an overall good quality and significant benefits were gained from optimising the procedure of the raw assembly. Among the parameters, in a transcriptome context, the N50 seems to be inappropriate due to the high variability of the transcriptome (i.e. very lowly vs very highly expressed transcripts) and because longer contigs do not necessarily reflect better assemblies (Salzberg et al. 2012). Accordingly, the N50 classical metric was also integrated with expression values to produce the Ex90N50, a more appropriate and biologically relevant metric for transcriptomes, although in our case the two results were in agreement. Furthermore, the redundancy of the transcriptome, which is expected with this kind of approach, was reduced following the optimisation procedure, as indicated by the decrease of complete and duplicated orthologues. This can be of particular interest in light of the effects that redundant transcripts can have on transcript quantification (Hsieh et al. 2019).

The tendency of some codons to be unevenly used in a transcriptome, guided by codon optimality, is a widespread process that holds from bacteria to higher eukaryotes (Hanson and Coller 2018). Codon usage preferences are indeed linked to mRNA stability, protein folding, translation efficiency and elongation, with highly expressed genes showing similar bias in codon content (reviewed in Chaney and Clark, 2015; Hanson and Coller, 2018; Novoa and Ribas de Pouplana, 2012). Genes enriched in optimal codons conserved between frog, fly, mouse and zebrafish and responsible for regulating mRNA stability were enriched in GO terms belonging to housekeeping processes (i.e. translation, protein folding, modulation of actin dynamics) during the maternal-to-zygotic transition (MZT) (Bazzini et al. 2016). Our results showed that more biased transcripts are mainly involved into the translational molecular machinery, ATP hydrolysis, muscle development and, remarkably, over-represented translation-related biological processes in the present study (see Supplementary file S4) matched exactly 4 out of 5 conserved processes between the aforementioned species for genes enriched in optimal codons. In the European sea bass, muscle development, protein biosynthesis and energy pathways were found to be, among others, differentially expressed during the larval ontogeny (Darias et al. 2008) and muscle development was specifically up-regulated in Atlantic bonito larvae at 30 dph (Sarropoulou et al. 2014). Similar mechanisms might be at play during the larval development of the Atlantic bluefin tuna, with more bias linked to important processes during this key period of development to meet the high demand for protein synthesis. Besides translation, several terms were also related to muscle development (striated muscle differentiation and development, sarcomere) and if biases found for these processes would contribute to the exceptional larval growth rate of this species still remains an open question. A similar point was addressed by Vicario et al. (2008), in which the very rapid stage-specific growth of Drosophila larvae was attributed to a global fast and accurate translation of expressed genes, which displayed the highest codon usage bias when compared with the embryo, adult and pupae stages. Nevertheless, our data was generated from a pool of several larval developmental stages, a point representing a potential confounding effect. Therefore, investigating separately these dynamics in different developmental stages will be essential to dig further into this scenario and understand how they are differentially modulated.

The majority of the protein-coding transcripts were functionally annotated, of which almost 80% were included in orthologous groups with other fish species. Intriguingly, some of the “assembly-specific” transcripts belonged to biological processes of great importance during the larval development such as protein digestion and absorption and iris morphogenesis. A precocious development of the digestive machinery was indeed highlighted during the Atlantic bluefin tuna larval development (Yúfera et al. 2014; Mazurais et al. 2015), raising the question of how evolution shaped these processes to sustain early piscivory. Greater activity levels of digestive enzymes were also found in the *Thunnus albacares* when compared with marine species of similar age/size (Buentello et al. 2011), supporting the concept of an adaptation of scombrid fish larvae to early piscivorous habits and high growth rates (Kaji et al. 2002). On the other side, the Pacific bluefin tuna *Thunnus orientalis* genome assembly allowed the discovery of evolutionary changes in several visual pigment genes which may have contributed to detecting contrasts in the blue-green spectrum and enhance hunting in the pelagic ocean (Nakamura et al. 2013). Recently, the effects of alternating light regimes on Atlantic bluefin tuna larval growth and survival rates were tested with promising results, highlighting an alternative solution for intensive cultures (Blanco et al. 2017). However, the authors also pointed out the possible existence of stage-dependent light needs and endocrine factors affecting appetite, digestion and growth during the larval development. To address such questions, our transcriptome comes at hand since provides a great amount of information at molecular level useful to investigate such processes in different larval stages and indicates that digestive capacity and eye morphogenesis might be interesting topics to be explored in greater details, as emphasised in other tuna species and also pointed out by Yúfera et al. (2014) for the Atlantic bluefin tuna.

In conclusion, in years of active interest towards the domestication of the Atlantic bluefin tuna, this work represents the first available de novo transcriptome assembly of the larval stages, a key point to close the life cycle of the species. Codon usage bias was identified for transcripts involved into translation and muscle development, a finding which supports the idea of codon preferences in biological processes needed to meet the high growth rate and demand for protein synthesis during the larval stages. Investigating these processes in different stages will therefore shed light on how they are regulated during the larval ontogeny. Future studies will surely benefit from the findings of our work due to the possibility to explore physiological responses at a molecular level, an important step to obtain a more complete picture regarding the biology of any species. Overall, this work represents a significant contribution to the molecular resources available for the Atlantic bluefin tuna.

## 5 Acknowledgements

The authors wish to truly thank Dr. Riccardo Aiese Cigliano for his support during the preparation of the present manuscript. Furthermore, the authors express their gratitude to Dr. Elena Sarropoulou and Dr. Giampaolo Zampicinini for sharing valuable information about the dataset.

## Bibliography

Afgan E, Baker D, Batut B, van den Beek M, Bouvier D, Čech M, Chilton J, Clements D, Coraor N, Grüning BA, Guerler A, Hillman-Jackson J, Hiltemann S, Jalili V, Rasche H, Soranzo N, Goecks J, Taylor J, Nekrutenko A, Blankenberg D (2018) The Galaxy platform for accessible, reproducible and collaborative biomedical analyses: 2018 update. Nucleic Acids Res 46:W537–W544. https://doi.org/10.1093/nar/gky379

Bazzini AA, del Viso F, Moreno-Mateos MA, Johnstone TG, Vejnar CE, Qin Y, Yao J, Khokha MK, Giraldez AJ (2016) Codon identity regulates mRNA stability and translation efficiency during the maternal-to-zygotic transition. EMBO J 35:2087–2103. https://doi.org/10.15252/embj.201694699

Benjamini Y, Hochberg Y (1995) Controlling the False Discovery Rate: A Practical and Powerful Approach to Multiple Testing. J R Stat Soc Ser B 57:289–300. https://doi.org/10.1111/j.2517-6161.1995.tb02031.x

Benzekri H, Armesto P, Cousin X, Rovira M, Crespo D, Merlo MA, Mazurais D, Bautista R, Guerrero-Fernández D, Fernandez-Pozo N, Ponce M, Infante C, Zambonino JL, Nidelet S, Gut M, Rebordinos L, Planas J V, Bégout M-L, Claros MG, Manchado M (2014) De novo assembly, characterization and functional annotation of Senegalese sole (Solea senegalensis) and common sole (Solea solea) transcriptomes: integration in a database and design of a microarray. BMC Genomics 15:952. https://doi.org/10.1186/1471-2164-15-952

Betancor MB, Ortega A, de la Gándara F, Tocher DR, Mourente G (2017) Lipid metabolism-related gene expression pattern of Atlantic bluefin tuna (Thunnus thynnus L.) larvae fed on live prey. Fish Physiol Biochem 43:493–516. https://doi.org/10.1007/s10695-016-0305-4

Betancor MB, Ortega A, de la Gándara F, Varela JL, Tocher DR, Mourente G (2019) Evaluation of different feeding protocols for larvae of Atlantic bluefin tuna (Thunnus thynnus L.). Aquaculture 505:523–538. https://doi.org/10.1016/J.AQUACULTURE.2019.02.063

Betancor MB, Sprague M, Ortega A, de la Gándara F, Tocher DR, Ruth R, Perkins E, Mourente G (2020) Central and peripheral clocks in Atlantic bluefin tuna(Thunnus thynnus, L.): Daily rhythmicity of hepatic lipid metabolism and digestive genes. Aquaculture 523:735220. https://doi.org/10.1016/j.aquaculture.2020.735220

Blanco E, Reglero P, Ortega A, de la Gándara F, Fiksen Ø, Folkvord A (2017) The effects of light, darkness and intermittent feeding on the growth and survival of reared Atlantic bonito and Atlantic bluefin tuna larvae. Aquaculture 479:233–239. https://doi.org/10.1016/j.aquaculture.2017.05.020

Bolger AM, Lohse M, Usadel B (2014) Trimmomatic: a flexible trimmer for Illumina sequence data. Bioinformatics 30:2114–2120. https://doi.org/10.1093/bioinformatics/btu170

Bourret J, Alizon S, Bravo IG (2019) COUSIN (COdon Usage Similarity INdex): A Normalized Measure of Codon Usage Preferences. Genome Biol Evol 11:3523–3528. https://doi.org/10.1093/gbe/evz262

Bray NL, Pimentel H, Melsted P, Pachter L (2016) Near-optimal probabilistic RNA-seq quantification. Nat Biotechnol 34:525–527. https://doi.org/10.1038/nbt.3519

Buentello JA, Pohlenz C, Margulies D, Scholey VP, Wexler JB, Tovar-Ramírez D, Neill WH, Hinojosa-Baltazar P, Gatlin Delbert. M. (2011) A preliminary study of digestive enzyme activities and amino acid composition of early juvenile yellowfin tuna (Thunnus albacares). Aquaculture 312:205–211. https://doi.org/10.1016/j.aquaculture.2010.12.027

Calduch-Giner JA, Bermejo-Nogales A, Benedito-Palos L, Estensoro I, Ballester-Lozano G, Sitjà-Bobadilla A, Pérez-Sánchez J (2013) Deep sequencing for de novo construction of a marine fish (Sparus aurata) transcriptome database with a large coverage of protein-coding transcripts. BMC Genomics 14:178. https://doi.org/10.1186/1471-2164-14-178

Carnevali O, Maradonna F, Sagrati A, Candelma M, Lombardo F, Pignalosa P, Bonfanti E, Nocillado J, Palma P, Gioacchini G, Elizur A (2019) Insights on the seasonal variations of reproductive features in the Eastern Atlantic Bluefin Tuna. Gen Comp Endocrinol 282:113216. https://doi.org/10.1016/j.ygcen.2019.113216

Carruthers M, Yurchenko AA, Augley JJ, Adams CE, Herzyk P, Elmer KR (2018) De novo transcriptome assembly, annotation and comparison of four ecological and evolutionary model salmonid fish species. BMC Genomics 19:32. https://doi.org/10.1186/s12864-017-4379-x

Cerdà J, Manchado M (2013) Advances in genomics for flatfish aquaculture. Genes Nutr 8:5–17. https://doi.org/10.1007/s12263-012-0312-8

Chaney JL, Clark PL (2015) Roles for Synonymous Codon Usage in Protein Biogenesis. Annu Rev Biophys 44:143–166. https://doi.org/10.1146/annurev-biophys-060414-034333

Chini V, Cattaneo AG, Rossi F, Bernardini G, Terova G, Saroglia M, Gornati R (2008) Genes expressed in Blue Fin Tuna (Thunnus thynnus) liver and gonads. Gene 410:207–213. https://doi.org/10.1016/j.gene.2007.12.012

Conesa A, Madrigal P, Tarazona S, Gomez-Cabrero D, Cervera A, McPherson A, Szcześniak MW, Gaffney DJ, Elo LL, Zhang X, Mortazavi A (2016) A survey of best practices for RNA-seq data analysis. Genome Biol 17:13. https://doi.org/10.1186/s13059-016-0881-8

Conway JR, Lex A, Gehlenborg N (2017) UpSetR: an R package for the visualization of intersecting sets and their properties. Bioinformatics 33:2938–2940. https://doi.org/10.1093/bioinformatics/btx364

Darias MJ, Zambonino-Infante JL, Hugot K, Cahu CL, Mazurais D (2008) Gene Expression Patterns During the Larval Development of European Sea Bass (Dicentrarchus Labrax) by Microarray Analysis. Mar Biotechnol 10:416–428. https://doi.org/10.1007/s10126-007-9078-1

De Metrio G, Bridges CR, Mylonas CC, Caggiano M, Deflorio M, Santamaria N, Zupa R, Pousis C, Vassallo-Agius R, Gordin H, Corriero A (2010) Spawning induction and large-scale collection of fertilized eggs in captive Atlantic bluefin tuna (Thunnus thynnus L.) and the first larval rearing efforts. J Appl Ichthyol 26:596–599. https://doi.org/10.1111/j.1439-0426.2010.01475.x

Delhomme N, Mähler N, Schiffthaler B, Sundell D, Mannepperuma C, Hvidsten TR (2014) Guidelines for RNA-Seq data analysis. Epigenesys Protoc 67:1–24

Emms DM, Kelly S (2015) OrthoFinder: solving fundamental biases in whole genome comparisons dramatically improves orthogroup inference accuracy. Genome Biol 16:157. https://doi.org/10.1186/s13059-015-0721-2

Emms DM, Kelly S (2019) OrthoFinder: phylogenetic orthology inference for comparative genomics. Genome Biol 20:238. https://doi.org/10.1186/s13059-019-1832-y

Fu L, Niu B, Zhu Z, Wu S, Li W (2012) CD-HIT: accelerated for clustering the next-generation sequencing data. Bioinformatics 28:3150–3152. https://doi.org/10.1093/bioinformatics/bts565

Gisbert E, Ortiz-Delgado JB, Sarasquete E (2008) Nutritional cellular biomarkers in early life stages of fish. Histol. Histopathol 23:1525–1539. https://doi.org/10.14670/HH-23.1525

Haas BJ, Papanicolaou A, Yassour M, Grabherr M, Blood PD, Bowden J, Couger MB, Eccles D, Li B, Lieber M, MacManes MD, Ott M, Orvis J, Pochet N, Strozzi F, Weeks N, Westerman R, William T, Dewey CN, Henschel R, LeDuc RD, Friedman N, Regev A (2013) De novo transcript sequence reconstruction from RNA-seq using the Trinity platform for reference generation and analysis. Nat Protoc 8:1494–1512. https://doi.org/10.1038/nprot.2013.084

Hanson G, Coller J (2018) Codon optimality, bias and usage in translation and mRNA decay. Nat Rev Mol Cell Biol 19:20–30. https://doi.org/10.1038/nrm.2017.91

Hara Y, Tatsumi K, Yoshida M, Kajikawa E, Kiyonari H, Kuraku S (2015) Optimizing and benchmarking de novo transcriptome sequencing: From library preparation to assembly evaluation. BMC Genomics 16:977. https://doi.org/10.1186/s12864-015-2007-1

Hsieh P-H, Oyang Y-J, Chen C-Y (2019) Effect of de novo transcriptome assembly on transcript quantification. Sci Rep 9:8304. https://doi.org/10.1038/s41598-019-44499-3

Huerta-Cepas J, Forslund K, Coelho LP, Szklarczyk D, Jensen LJ, von Mering C, Bork P (2017) Fast Genome-Wide Functional Annotation through Orthology Assignment by eggNOG-Mapper. Mol Biol Evol 34:2115–2122. https://doi.org/10.1093/molbev/msx148

Hughes LC, Ortí G, Huang Y, Sun Y, Baldwin CC, Thompson AW, Arcila D, Betancur-R. R, Li C, Becker L, Bellora N, Zhao X, Li X, Wang M, Fang C, Xie B, Zhou Z, Huang H, Chen S, Venkatesh B, Shi Q (2018) Comprehensive phylogeny of ray-finned fishes (Actinopterygii) based on transcriptomic and genomic data. Proc Natl Acad Sci 115:6249–6254. https://doi.org/10.1073/pnas.1719358115

ICCAT (2008) Recommendaton amending the recommendation by ICCAT to establish a multiannual recovery plan for bluefin tuna in the eastern Atlantic and Mediterranean. Madrid

Kaji T, Kodama M, Arai H, Tagawa M, Tanaka M (2002) Precocious development of the digestive system in relation to early appearance of piscivory in striped bonito Sarda orientalis larvae. Fish Sci 68:1212–1218. https://doi.org/10.1046/j.1444-2906.2002.00557.x

Kassambara A (2020) ggpubr: “ggplot2” Based Publication Ready Plots

Kopylova E, Noé L, Touzet H (2012) SortMeRNA: fast and accurate filtering of ribosomal RNAs in metatranscriptomic data. Bioinformatics 28:3211–3217. https://doi.org/10.1093/bioinformatics/bts611

Langmead B, Salzberg SL (2012) Fast gapped-read alignment with Bowtie 2. Nat Methods 9:357–359. https://doi.org/10.1038/nmeth.1923

Letunic I, Bork P (2007) Interactive Tree Of Life (iTOL): an online tool for phylogenetic tree display and annotation. Bioinformatics 23:127–128. https://doi.org/10.1093/bioinformatics/btl529

Louro B, Passos ALS, Souche EL, Tsigenopoulos C, Beck A, Lagnel J, Bonhomme F, Cancela L, Cerdà J, Clark MS, Lubzens E, Magoulas A, Planas J V, Volckaert FAM, Reinhardt R, Canario AVM (2010) Gilthead sea bream (Sparus auratus) and European sea bass (Dicentrarchus labrax) expressed sequence tags: Characterization, tissue-specific expression and gene markers. Mar Genomics 3:179–191. https://doi.org/10.1016/j.margen.2010.09.005

MacKenzie BR, Mosegaard H, Rosenberg AA (2009) Impending collapse of bluefin tuna in the northeast Atlantic and Mediterranean. Conserv Lett 2:26–35. https://doi.org/10.1111/j.1755-263X.2008.00039.x

MacManes MD (2014) On the optimal trimming of high-throughput mRNA sequence data. Front Genet 5:13. https://doi.org/10.3389/fgene.2014.00013

Mazurais D, Covès D, Papandroulakis N, Ortega A, Desbruyeres E, Huelvan C, Le Gall MM, de la Gándara F, Cahu CL (2015) Gene expression pattern of digestive and antioxidant enzymes during the larval development of reared Atlantic bluefin tuna (ABFT), Thunnus thynnus L. Aquac Res 46:2323–2331. https://doi.org/10.1111/are.12387

McMenamin SK, Parichy DM (2013) Chapter Five - Metamorphosis in Teleosts. In: Shi Y-BBT-CT in DB (ed) Animal Metamorphosis. Academic Press, pp 127–165

Medina A, Aranda G, Gherardi S, Santos A, Mèlich B, Lara M (2016) Assessment of spawning of Atlantic bluefin tuna farmed in the western Mediterranean Sea. Aquac Environ Interact 8:89–98. https://doi.org/10.3354/aei00166

Moreton J, Izquierdo A, Emes RD (2016) Assembly, Assessment, and Availability of De novo Generated Eukaryotic Transcriptomes. Front. Genet. 6:361. https://doi.org/10.3389/fgene.2015.00361

Moriya Y, Itoh M, Okuda S, Yoshizawa AC, Kanehisa M (2007) KAAS: an automatic genome annotation and pathway reconstruction server. Nucleic Acids Res 35:W182–W185. https://doi.org/10.1093/nar/gkm321

Mylonas CC, Bridges C, Gordin H, Ríos AB, García A, De La Gándara F, Fauvel C, Suquet M, Medina A, Papadaki M, Heinisch G, De Metrio G, Corriero A, Vassallo-Agius R, Guzmán JM, Mañanos E, Zohar Y (2007) Preparation and administration of gonadotropin-releasing hormone agonist (GnRHa) implants for the artificial control of reproductive maturation in captive-reared Atlantic bluefin tuna (Thunnus thynnus thynnus). Rev Fish Sci 15:183–210. https://doi.org/10.1080/10641260701484572

Mylonas CC, de la Gándara F, Corriero A, Ríos AB (2010) Atlantic bluefin tuna (thunnus thynnus) farming and fattening in the mediterranean sea. Rev Fish Sci 18:266–280. https://doi.org/10.1080/10641262.2010.509520

Nakamura Y, Mori K, Saitoh K, Oshima K, Mekuchi M, Sugaya T, Shigenobu Y, Ojima N, Muta S, Fujiwara A, Yasuike M, Oohara I, Hirakawa H, Chowdhury VS, Kobayashi T, Nakajima K, Sano M, Wada T, Tashiro K, Ikeo K, Hattori M, Kuhara S, Gojobori T, Inouye K (2013) Evolutionary changes of multiple visual pigment genes in the complete genome of Pacific bluefin tuna. Proc Natl Acad Sci 110:11061–11066. https://doi.org/10.1073/pnas.1302051110

Nishimura O, Hara Y, Kuraku S (2017) gVolante for standardizing completeness assessment of genome and transcriptome assemblies. Bioinformatics 33:3635–3637. https://doi.org/10.1093/bioinformatics/btx445

Novoa EM, Ribas de Pouplana L (2012) Speeding with control: codon usage, tRNAs, and ribosomes. Trends Genet 28:574–581. https://doi.org/10.1016/j.tig.2012.07.006

Pauletto M, Manousaki T, Ferraresso S, Babbucci M, Tsakogiannis A, Louro B, Vitulo N, Quoc VH, Carraro R, Bertotto D, Franch R, Maroso F, Aslam ML, Sonesson AK, Simionati B, Malacrida G, Cestaro A, Caberlotto S, Sarropoulou E, Mylonas CC, Power DM, Patarnello T, Canario AVM, Tsigenopoulos C, Bargelloni L (2018) Genomic analysis of Sparus aurata reveals the evolutionary dynamics of sexbiased genes in a sequential hermaphrodite fish. Commun Biol 1:119. https://doi.org/10.1038/s42003-018-0122-7

Reglero P, Ortega A, Blanco E, Fiksen Ø, Viguri FJ, de la Gándara F, Seoka M, Folkvord A (2014) Size-related differences in growth and survival in piscivorous fish larvae fed different prey types. Aquaculture 433:94–101. https://doi.org/10.1016/j.aquaculture.2014.05.050

Rønnestad I, Yúfera M, Ueberschär B, Ribeiro L, Sæle Ø, Boglione C (2013) Feeding behaviour and digestive physiology in larval fish: current knowledge, and gaps and bottlenecks in research. Rev Aquac 5:S59–S98. https://doi.org/10.1111/raq.12010

RStudio team (2015) RStudio: Integrated Development for R

Salzberg SL, Phillippy AM, Zimin A, Puiu D, Magoc T, Koren S, Treangen TJ, Schatz MC, Delcher AL, Roberts M, Marçais G, Pop M, Yorke JA (2012) GAGE: A critical evaluation of genome assemblies and assembly algorithms. Genome Res 22:557–567. https://doi.org/10.1101/gr.131383.111

Sarropoulou E, Moghadam HK, Papandroulakis N, De la Gándara F, Ortega Garcia A, Makridis P (2014) The Atlantic Bonito (Sarda sarda, Bloch 1793) Transcriptome and Detection of Differential Expression during Larvae Development. PLoS One 9:e87744. https://doi.org/10.1371/journal.pone.0087744

Sawada Y, Okada T, Miyashita S, Murata O, Kumai H (2005) Completion of the Pacific bluefin tuna Thunnus orientalis (Temminck et Schlegel) life cycle. Aquac Res 36:413–421. https://doi.org/10.1111/j.1365-2109.2005.01222.x

Schmieder R, Edwards R (2011) Quality control and preprocessing of metagenomic datasets. Bioinformatics 27:863–864. https://doi.org/10.1093/bioinformatics/btr026

Simão FA, Waterhouse RM, Ioannidis P, Kriventseva E V., Zdobnov EM (2015) BUSCO: assessing genome assembly and annotation completeness with single-copy orthologs. Bioinformatics 31:3210–3212. https://doi.org/10.1093/bioinformatics/btv351

Slowikowski K (2020) ggrepel: Automatically Position Non-Overlapping Text Labels with ggplot2

Smith-Unna R, Boursnell C, Patro R, Hibberd JM, Kelly S (2016) TransRate: reference-free quality assessment of de novo transcriptome assemblies. Genome Res 26:1134–1144. https://doi.org/10.1101/gr.196469.115

Suda A, Nishiki I, Iwasaki Y, Matsuura A, Akita T, Suzuki N, Fujiwara A (2019) Improvement of the Pacific bluefin tuna (Thunnus orientalis) reference genome and development of male-specific DNA markers. Sci Rep 9:14450. https://doi.org/10.1038/s41598-019-50978-4

Supek F, Bošnjak M, Škunca N, Šmuc T (2011) REVIGO Summarizes and Visualizes Long Lists of Gene Ontology Terms. PLoS One 6:e21800. https://doi.org/10.1371/journal.pone.0021800

Tenenbaum D (2018) KEGGREST: Client-side REST access to KEGG

Tine M, Kuhl H, Gagnaire P-A, Louro B, Desmarais E, Martins RST, Hecht J, Knaust F, Belkhir K, Klages S, Dieterich R, Stueber K, Piferrer F, Guinand B, Bierne N, Volckaert FAM, Bargelloni L, Power DM, Bonhomme F, Canario AVM, Reinhardt R (2014) European sea bass genome and its variation provide insights into adaptation to euryhalinity and speciation. Nat Commun 5:5770. https://doi.org/10.1038/ncomms6770

Trumbić Ž, Bekaert M, Taggart JB, Bron JE, Gharbi K, Mladineo I (2015) Development and validation of a mixed-tissue oligonucleotide DNA microarray for Atlantic bluefin tuna, Thunnus thynnus (Linnaeus, 1758). BMC Genomics 16:1007. https://doi.org/10.1186/s12864-015-2208-7

Vicario S, Mason CE, White KP, Powell JR (2008) Developmental Stage and Level of Codon Usage Bias in Drosophila. Mol Biol Evol 25:2269–2277. https://doi.org/10.1093/molbev/msn189

Wickham H (2016) ggplot2: Elegant Graphics for Data Analysis. Springer-Verlag New York

Wood DE, Lu J, Langmead B (2019) Improved metagenomic analysis with Kraken 2. Genome Biol 20:257. https://doi.org/10.1186/s13059-019-1891-0

Wright F (1990) The ‘effective number of codons’ used in a gene. Gene 87:23–29. https://doi.org/10.1016/0378-1119(90)90491-9

Yu G, Wang L-G, Han Y, He Q-Y (2012) clusterProfiler: an R Package for Comparing Biological Themes Among Gene Clusters. Omi A J Integr Biol 16:284–287. https://doi.org/10.1089/omi.2011.0118

Yúfera M, Ortiz-Delgado JB, Hoffman T, Siguero I, Urup B, Sarasquete C (2014) Organogenesis of digestive system, visual system and other structures in Atlantic bluefin tuna (Thunnus thynnus) larvae reared with copepods in mesocosm system. Aquaculture 426–427:126–137. https://doi.org/10.1016/J.AQUACULTURE.2014.01.031

Zhi-Liang H, Bao J, James M. R (2008) CateGOrizer: A web-based program to batch gene ontology classification categories. Online J Bioinforma 9:108–112

Zohar Y, Mylonas CC, Rosenfeld H, de la Gándara F, Corriero A (2016) Chapter 7 - Reproduction, Broodstock Management, and Spawning in Captive Atlantic Bluefin Tuna. In: Benetti DD, Partridge GJ, Buentello ABT-A in TA (eds) Advances in Tuna Aquaculture. Academic Press, San Diego, pp 159–188

